# Viscoelastic flow-integrated Raman spectroscopy platform^†^

**DOI:** 10.64898/2026.01.13.699233

**Authors:** Adam Stovicek, Murat Serhatlioglu, Babak Rezaei, Stephan Sylvest Keller, Barth F. Smets, Mikkel Bentzon-Tilia, Anders Kristensen, Arnaud Dechesne

## Abstract

We demonstrate the first viscoelastic-based flow-integrated Raman spectroscopy (FIRS) platform using spontaneous Raman spectroscopy with a simplified fluidic design that precisely controls particle transit speed and predicts arrival time to the Raman interrogation region, together with time synchronized triggered acquisition. Using viscoelastic flow focusing and optical detection of velocity for hardware triggering, the setup enables high-throughput, real-time Raman analysis of particles in continuous flow. The system operates with a single pressure pump, two sets of fiber-coupled lasers and detectors, and a microcapillary precisely aligned in a projection micro-stereolithography 3D-printed mount, and ensures detection stability across various flow rates. The demonstrated platform is modular and scalable. Detection rates of up to 80 cells per minute were achieved and dinoflagellate *Alexandrium ostenfeldii* cultures were analyzed. The integrated optical trigger mechanism supports future extensions such as cell sorting or additional analyses. A user-friendly graphical interface provides full control of fluidics, spectroscopy, and real-time monitoring, making this compact, low-cost FIRS platform accessible across a wide range of laboratory environments.

## 1. Introduction

Phenotypic profiling of single cell populations or cell clusters is a valuable tool in microbial ecology, industrial fermentation bio-engineering and biomedicine. Morphology based phenotypic profiling can for example identify human cell early apoptosis ^1^, determine the physiological impact of secondary metabolites, aiding in the search of new therapeutics ^2^, reveal circulating tumor cells in blood ^3^, or assess antibiotic resistant subpopulations in cultures of *Staphylococcus aureus* ^4^. Such profiling approaches are often based on image analysis of multiple overlayed fluorescence channels. However, this approach is inherently confined to a limite set of targeted fluorescent probes.

Raman spectroscopy (RS) is established as a label free analytical tool providing robust information on biological samples suspended in aqueous buffer ^5^. RS enables, for example, phenotypic profiling of bacteria resistant to antimicrobial compounds ^6^, surveying yeast phenotypes (e.g. carotene production or physiological impact of ethanol) ^7,8^, or mapping algae phenotypic diversity ^9^. Furthermore, RS can follow the intake of stable isotopes of carbon, nitrogen or deuterium ^10,11^, thus describing the profile of metabolic activities within a cell population. Several flow based systems incorporating Raman detectors have been developed to profile metabolic activity of cell populations ^12^ or to assess secondary metabolite production ^13^.

Depending on the excitation scheme, Raman spectroscopy can be implemented in various forms, including spontaneous, resonance, and coherent techniques ^14^. A main drawback of spontaneous RS is characterized by a low signal yield, typically requiring exposure times of at least 10s of ms. This limits the throughput of any flow based method. Currently published high throughput methods are enabled by enhanced Raman scattering yield due to resonant Raman spectra ^13,15^ (up to 2700 cells/min ^13^) or by recording only specific spectral bands ^16^ (300 cells/min). Raman scattering can be increased using coherent Raman spectroscopy, thus enabling higher throughput (3000 cells/min ^17^). However this method currently only provides lower spectral resolution compared to spontaneous Raman spectroscopy ^18,19^ and adds substantial system development cost. There is currently an inherent tradeoff between data acquisition speed and signal resolution.

The requirement for an extended exposure time can be accommodated by incorporating cell immobilization during exposure. This often requires relatively complex microfluidic setups, fabricated from polymers, such as polydimethylsiloxane (PDMS) ^15,16,20^. These polymers generate a strong Raman background and only allow detection of resonance Raman spectra of e.g. carotenoids ^15^. In contrast, dielectrophoretic (DEP) based capturing methods have been implemented on quartz chips ^13^. However, these systems support only low conductivity buffers. Higher conductivity buffers induce power loss due to Joule heating, causing sample damage ^21^. Consequently, this method is incompatible with high salinity biological samples (e.g. marine samples) due to osmotic pressure induced lysis.

In contrast to immobilization based methods, spontaneous Raman acquisition on free flowing samples requires a precise synchronization with the particle arrival. Precise synchronization is therefore a primary constraint for acquiring single-particle Raman spectra in flow ^22^. This is generally achieved by a set of sensors that detect incoming particles and trigger a downstream analytical device such as a Raman detector or a particle sorting device ^15–17^. Most of these designs are built around a single computer with Windows OS running LabVIEW (National Instruments) or an equivalent software together with data acquisition hardware ^15,17^ . This design limits tasks requiring precision scheduling of an event downstream of a detector, due to Windows OS’ inherent scheduling latency, which was reported in order of 100s of ms ^25^. This calls for alternative solutions for time sensitive scheduling applications.

Particles in any type of flow system need to be deterministically located to be accessible to the narrow detection voxel of a Raman excitation laser. This alignment can be achieved by DEP focusing ^13^, or electroacoustics focusing ^17,26^. Many previous Raman flow applications rely on hydrodynamic focusing ^16,20-27-29^ . However, sheath flow hydrodynamics and pump fluctuations may introduce velocity jitter, inflating coefficients of variation and destabilizing particle position at the probe especially at high flow rates. These effects impose increasing challenges at higher sample pressure and flow rate ^30,31^.

Viscoelastic microfluidics ^32,33^ provides a fluidic solution to reduce timing uncertainty. In dilute viscoelastic polymer solutions, elastic normal stresses generate lateral lift forces that drive particles to a narrow centerline equilibrium in straight channels without sheath flow in a single inlet/outlet configuration. The viscoelastic flow focusing strength depends on channel geometry, the polymer relaxation time and Weissenberg number. Stable 3D viscoelastic flow focusing has been shown over a wide range of flow conditions with straight microfluidic channels ^34–36^. Reviews and experimental studies in glass and polymer microchannels, including viscoelastic solutions with hyaluronic acid (HA), confirm that viscoelastic focusing reduces lateral wander and axial velocity dispersion at the interrogation point ^32,37-39^ . By transforming a stochastic arrival process into a predictable one, viscoelastic focusing enables the use of narrow acquisition windows.

Here, we present a sheathless, time-triggered Raman flow cytometer in which synchronization is the primary design driver. A straight circular glass microcapillary with 250 µm inner diameter which carries particles suspended in a HA solution, where they are centered by elastic lift, enabling reproducible acquisition of single-particle Raman spectra without the complexity of sheath flow. A 3D-printed mount secures the sample capillary and two orthogonal fibers which form two forward scatter detectors used to measure transit time and velocity. The speed of each particle is calculated separately, further accounting for temporal variation in the flow. This speed information is then used as a trigger signal for downstream in-line events (e.g. cell sorting or Raman acquisition). The system implements low latency detection and a trigger layer comprised of cheap off-the-shelf Linux based components, together with a standard latency PC hosting the user interface and the computationally expensive tasks. Importantly, the modular, low-cost design lowers the technical barrier for adoption of Raman cytometry in diverse laboratory settings.

## 2 Materials and Methods

### 2.1 Alignment device fabrication

A 3D-printed device was engineered to align a fused silica capillary containing the viscoelastic solution with perpendicular optical fibers. This setup includes a perpendicular arrangement of collinear optical fibers forming two forward scatter detectors to detect incoming particles (Fig. 1), and an opposing set of optical fibers for collecting transmitted light. The device was designed using SolidWorks CAD modeling software (version 2023). Fabrication was performed with a projection microstereolithography (PµSLA) 3D printer (NanoArch S140, Boston Micro Fabrication Inc., USA), achieving an XY resolution of 10 µm and a Z resolution of 5 µm. The photocurable resin (HTL, BMF, USA) was cured using 405 nmlight, with an exposure time of 2 s for the initial 10 layers and 1 s for the subsequent layers. The light intensity was set to 20 % of maximum intensity for the first 10 layers and 10 % of maximum intensity 110 mW*/*cm^2^ for the remaining layers.

**Fig. 1.**
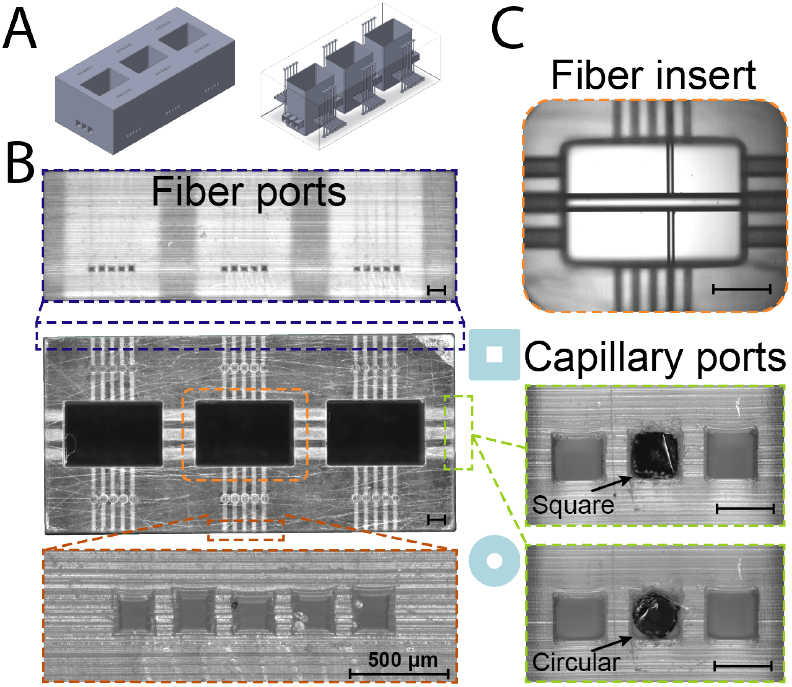
Overview of the 3D-printed fiber alignment device. (A) CAD drawing illustrates 3D structure of the device. (B) Microscope images showing the top-view and optical fiber ports in full scale, and side-view for fiber ports in close-up. 3 compartments are designed as forward scattering channels (only 2 of them used in this study). The alignment device is 12.5×6 *mm*^2^ and the rectangular windows are 3×2 *mm*^2^. All ports are designed rectangular. Optical fiber ports are designed in two different widths: 127 µm and 130 µm. Aspect ratio of 3 different values are preferred (0.67, 0.71, and 0.77) to compensate the sagging of the roof in fiber and capillary inserts. (C) Microcopy image of fiber and a circular optical capillary inserted in ch1 section. The optimum fiber and capillary fit was tested by inserting fibers and capillaries to their ports one-by-one and identifying the perfect fit from microscopic observations. The fused silica capillary ports of 400 µm, AR= 0.8, 0.77, and 0.74 tested with square and circular cross section capillaries and perfect fit was achieved in AR=1.30. Refer to Supplementary file for complete set of characterization steps for the 3D printed alignment device

Free surface PµSLA performed without sacrificial layers is prone to surface deformation of the liquid resin, which can propagate into the cured geometry and induce dimensional deviations in the fabricated structure ^40^. Printed features showed significant deviations from the designed dimensions for features like circle, square, and triangle, and thus required compensation of the projection pattern ^41^. In our designs, the 3D printed part consists of alignment guides for the fused silica capillary (Outer Diameter: 360 *±* 10 µm with approx. 18 µm polymer coating) and optical fibers (Cladding Diameter: 125 *±* 1 µm with Coating Diameter: 245 *±* 15 µm). The aim was to fabricate square cross-section alignment guides, fit to insert size; however, due to sagging of the free top-surface (roof) and material shrinkage inherent to the 3D printing process, it was not possible to directly obtain square geometries from the nominal designs. A systematic optimization procedure was therefore conducted to identify the design parameters that yield near-square cross-section guide geometries after fabrication. Please refer to Supplementary file for complete set of characterization steps for the 3D printed alignment device (see Fig. S1-S4).

### 2.2 Speed analysis setup

As particles pass through the forward scatter detectors they eclipse the incoming laser beam (wavelength 530 nm), resulting in a decrease in the light/voltage signal (Fig 2B). Particles first pass through optical channel 1 (ch1), located 4 mm upstream of optical channel 2 (ch2). The transmitted laser light was collected by an optical fiber, which directed the light to an optical detector. The resulting voltage signal was measured using a Raspberry Pi 5 (RPi5) equipped with a DAQ card (MCC128, Digilent, USA). This setup continuously acquired data from the forward scatter detectors at a rate of approximately 2 kHz. Each measurement point was individually time-stamped, and the data was aggregated into packets of 200 measurements. These data packets were submitted to a REDIS database, from which they were periodically retrieved by the central Windows system. The retrieved data was then analyzed and displayed for the user.

**Fig. 2.**
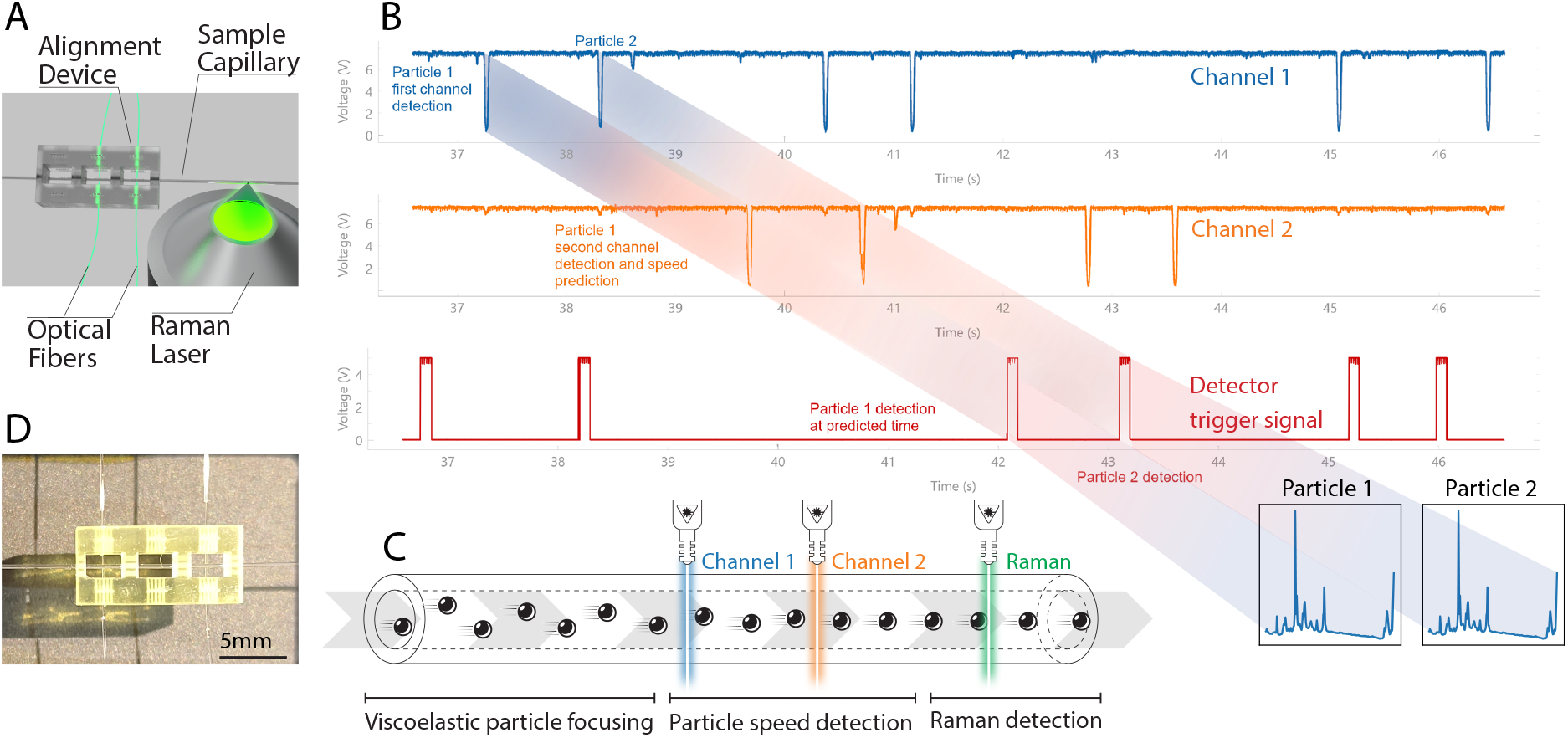
Schematic representation of the device design. (A) Particles flow from left to right through a fused-silica capillary, where they first encounter a set of lasers detecting their passage and speed (channel 1 and channel 2). This information is used to determine individual particle speed and calculate particle arrival time at the detector. (B) The 3D printed fiber alignment device with incoming optical fibers and colinear detection fibers. (C) The resulting data from channel 1 (blue) and 2 (orange) from the data acquisition device and the resulting trigger signal for the Raman detector (red). (D) The entire system is operated from a graphical interface.

### 2.3 Speed analysis algorithm

The passages of particles in front of the two forward scatter detectors were identified using the find_peaks method from the Python SciPy package (v1.11.3). The leading edge of each peak, referred to as the “left_ips” in find_peaks, was recorded as the peak position and further analyzed using a speed prediction algorithm (see below). The speed prediction begins by recording a peak timestamp *t*_*n,ch*2_ in ch2. The transit time Δ*t* estimate from ch1 to ch2 is then calculated using the nearest preceding peak in ch1. This initial estimate is validated by applying the same duration to a prior peak in ch2, as described by Equation 1:

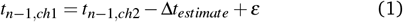

Where *ε* represents a user-defined tolerance error. The resulting value *t*_*n*−1,*ch*1_ is expected to closely match an existing peak in ch1 if Δ*t*_*estimate*_ is accurate. This process is iterated for a user-defined number of peaks (default is 5), and the results are validated against a user-defined threshold of successful matches (default is 3). If validation fails, the procedure is repeated using a preceding peak in ch1 to generate a new Δ*t*_*estimate*_. This iterative process continues for a user-defined number of attempts (default is 10). This method leverages the randomness of particle distribution in the flow and it assumes a low fluctuation of flow speed among nearby particles, however it is robust against speed fluctuations at longer timescales. Raw data from a data acquisition card can be visualized using a GUI interface [Daq Data Analyzer](https://github.com/AdamSorrel/daqDataAnalyzer) written for this project.

### 2.4 Raman spectroscopy setup

The fluidic setup is portable and can be set up on any inverted microscope, see photo in Fig 2B. The data was acquired on a microscope (Nikon Eclipse TE2000-U, Japan) equipped with a 532 nm continuous wave excitation laser source (Cobolt 08-01) focused with a 50x, 0.8 NA objective onto a fused silica capillary. This objective focuses the laser beam to a spot of 2.86 µm and depth of focus of 20 µm, measured using the razor-edge method ^42^. The resulting Stokes Raman scattered light is collected by the same objective and separated from the excitation laser by a dichroic mirror. The collected Rayleigh scattering is filtered by a long-pass edge filter (RazorEdge LP Edge Filter 532 RU, Semrock, USA) and a notch filter. The resulting Stokes Raman scattered light is analyzed using an imaging spectrograph (Shamrock SR-500i, Andor) with a 300 µm wide slit and an additional long pass filter. The wavelengths were separated using a 1200 lines/mm diffraction grating and the resulting spectrum was analyzed by a CCD camera (Newton 920, Andor). The read-out sensor was cooled to −70 ^◦^C to reduce the effect of dark current.

#### 2.4.1 System latency measurements

The latency of a native kernel of the Raspbian operating system (OS) was compared with that of partially and fully PREEMPT_RT patched OS versions. This comparison was conducted under stress conditions generated by the hackbench scheduler stress test and cyclictest, both of which are included in the rt-tests package v1.0. Detailed procedural information and all applied settings are provided in **Supplementary File 1**. In summary, stress was induced on each core by instructing it to send 50,000 messages of 100 bytes each. During this load, a cyclictest was performed to retrieve latency results.

The system latency was further measured using a square wave generator (HM 8030-S, Hameg, Germany). The resulting square wave was captured using the Raspbery Pi 5 associated DAQ (MCC 128, Digilent, USA). Measuring the adjacent quare waves yielded the error of the DAQ measurement.

### 2.5 False positive and negative quantification

False positive samples were quantified by checking the content of the spectra. Main PS peak was verified identifying benzene ring vibration around 561.75 nm (995 cm) and calculating difference with neighboring spectral region. These differences were subsequently plotted in a histogram and an appropriate cutoff was determined visually (Supplementary Fig. S5). False negative detections were inferred from particle detection in speed measuring laser one, which did not yield a spectrum acquisition event. The proportion of crowded particles that arrived to the detector within 200 ms, which is the limit of repeatability of measurement for the tested Raman camera, was calculated from time stamp differences between all particles as measured by channel 1 of the speed measuring laser.

### 2.6 Automation platform architecture

The presented automation platform provides a plug-in based platform operated by a Windows computer, which provides a high performance, but also a high latency time, asynchronous part of the system. This is combined with two RPi5 units, running a PREEMPT_RT Raspbian Linux kernel, providing a low performance and low latency time synchronous layer. A common time is established between the two RPi5 units using a Precision Time Protocol (PTP) implemented in ptp4l (3.1.1) and phc2sys (3.1.1) commands ^43^.

One RPi5 computer serves as the PTP grandmaster clock, the host of a noSQL REDIS database (version) and the Transistor-Transistor Logic (TTL) signal trigger for the Raman detector. The hosted REDIS database serves as a communication nexus between the asynchronous Windows computer and the two RPi5 devices. The other RPi5 unit is dedicated to data acquisition from the two laser channels for particle speed estimation. The time stamps corresponding to particle passage in front of the speed measurement lasers are then shared with the Windows computer through the REDIS database. The Windows computer then calculates the particle speed. The particle speed prediction ultimately produces a time stamp of a trigger signal corresponding to the expected time of particle passage in front of a detector. This signal is returned to a time synchronized RPi5 computer via REDIS database, which is continuously polled by a Python program. Upon receiving trigger time stamps, this program then schedules a trigger signal using Python sched module which is spawned as a new process to allow maximum OS timing flexibility.

### 2.7 Sample preparation

A viscoelastic solution of HA was prepared by dissolving 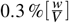 HA (1.0 - 2.0 MDa, Carbosynth, UK) in MilliQ filtered water. The solution was subjected to continuous mixing for 24 h using a magnetic stirrer. Post-mixing, the viscoelastic solution was stored at 4 ^◦^C until further use. Prior to each experimental run, the capillary of the device was rinsed sequentially with MilliQ filtered water and the 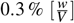 HA viscoelastic solution.

Spherical polystyrene (PS) beads (Thermofisher Scientific, USA) with 50 µm diameter were centrifuged (Eppendorf, 5430, Germany) for 5 min 500 g. The supernatant was discarded and replaced with the 2 % HA viscoelastic solution. The beads were maintained in suspension within an Eppendorf tube through continuous agitation with a magnetic stirrer.

Cells of the alga *Alexandrium ostenfeldii* were grown in f/2 media ^44^ at 19 ^◦^C with illumination strength 50 µmol*/*(m^2^ s). The resulting culture, containing both living and dead algal cells was centrifuged at 5000 g for 5 min and the supernatant was discarded. The pellet was fixed with 3 % formaldehyde and resuspended in a 3 % Instant Ocean (Aquarium Systems, Sarrebourg, France) solution. A 10 µL aliquot of the resulting culture was resuspended in 0.490 mL of 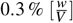 HA viscoelastic buffer solution prepared with 3 % Instant Ocean immediately before the experiment.

### 2.8 Flow conditions

The flow was performed in a 3000 ppm solution of HA in a capillary with internal diameter 250 µm. Algae cells or PS beads were mixed continuously with a magnetic stirrer. Particle speed was adjusted with a pressure pump (Fluigent MFSC, France) with pressure between 100 mbar and 900 mbar. The focusing performance of the particles was evaluated in this range prior to the experiments both at the entrance to the capillary and at the Raman acquisition zone. The particle position data was acquired using a high speed camera (AOS S-MOTION) recording at 500 FPS frame rate and the particle trajectories were generated by image stacking of frames using a Matlab script (MATLAB 2023).

## 3 Results and discussion

We present experiments validating the flow focusing performance, timing analysis of the device data acquisition and trigger and quantification of false positive and false negative events. The device maximum throughput was measured with PS beads. The ability to acquire biological data was validated using dinoflagellate alga *Alexandrium ostenfeldii*.

### 3.1 Flow focusing validation

Flow focusing performance for the particles at pressure ranges (100 mbar–900 mbar) was tested with a high speed camera. We observed a good particle alignment across the tested speeds with central deviation of 1.4 *±* 0.3 µm for 100 mbar pressure, −7.0 *±* 9.1 µm for 500 mbar and 0.8 *±* 0.4 µm for 900 mbar, which is an acceptable performance given the size of the particles see Supplementary Table S1.

### 3.2 System scheduling latency

The architecture for time synchronization is outlined in Fig. 3. To mitigate OS timing latency, we use a pseudo-real-time Raspbian OS patched with a PREEMPT_RT. We tested latency under high load of a native Raspbian OS, partially patched and fully patched system (Fig. S6), upon which we decided to use a fully patched system with the highest latency ≤ 50 µs (Fig. S6, C).

**Fig. 3.**
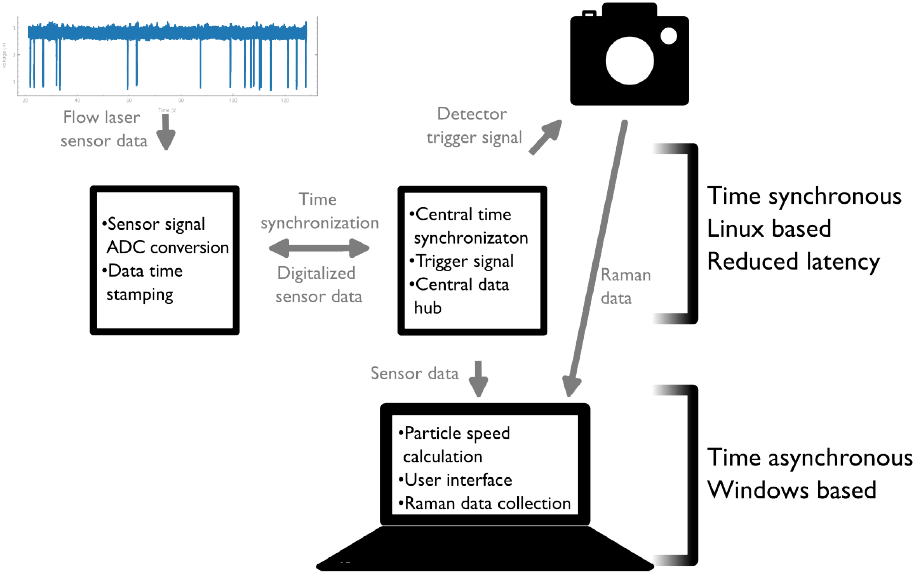
The automation platform comprises of a time synchronous setup of low latency Raspberry Pi devices, which are collecting the data from the speed measuring detection channels using an analog-digital converter (ADC). This data is time stamped and collected in a database running on a second low latency Raspberry Pi device, which is also running the trigger routine as well as a high precision PTP time synchronization server. The speed sensor data is collected in a Windows computer, running the user interface as well as drivers for the Raman detector. This central unit calculates particle speed and issues back a trigger time stamp which is collected by the Raspberry Pi.

**Fig. 4.**
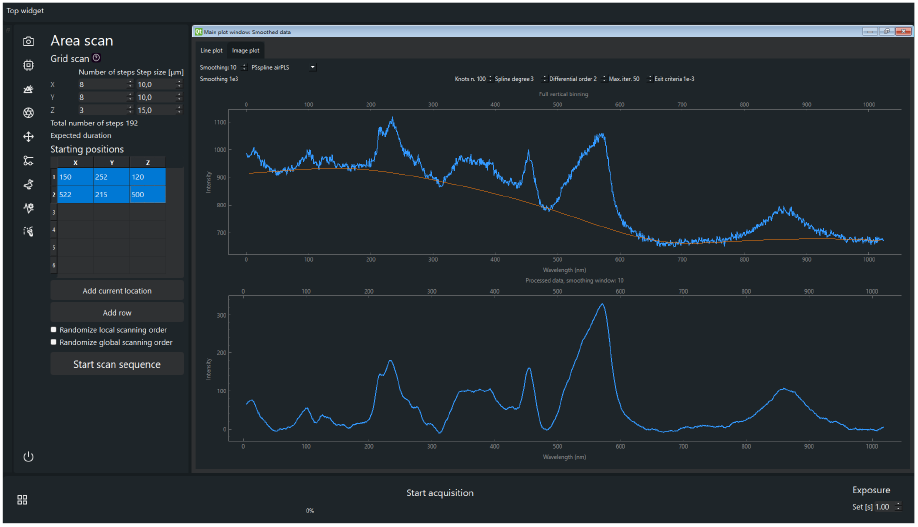
Graphical user interface of the central driving program with the individual plug-in extensions (left bar), settings area and a plot area in the middle with multiple user adjustable plot windows.

### 3.3 System timing error

In addition, we tested the DAQ card time stamping method as a possible source of error. Briefly, an externally generated square wave was captured with the DAQ card and the edge distances in the resulting time stamped data were compared. The dispersion of these values was interpreted as an error with a standard deviation of 0.66 ms and maximum extreme values of 1.98 ms and −1.69 ms respectively. The histogram of these errors is presented in Fig. S7.

The trigger signal can also act as a further source of timing error. To evaluate this, a desired trigger signal timestamp was compared to the actual trigger time stamp. The average value of the trigger error was −89.3 µs with standard deviation 60.8 µs (n = 986). The complete results are presented in Fig. S8.

### 3.4 Graphical user interface

The whole system is operated from a central program with a graphical user interface (Fig. 2D), allowing users to set operational parameters for the Raman detector, pressure flow, and particle detection data acquisition parameters including threshold peak size in both directions from the baseline and minimum peak distance. Furthermore, the system provides real time data processing such as Raman spectrum smoothing and baseline subtraction. Moreover, the system incorporates area scanning in a 3 coordinate system with the option to choose multiple starting positions for each scan. This platform is developed with a plug-in based architecture and extendability in mind with a single point of entry for new modules and their relatively simple integration.

### 3.5 Speed and position prediction testing with PS beads

We have tested the system performance with in-flow Raman detection of polystyrene beads. Fig. 5AC shows their representative spectra as well as the buffer background for each type of the sample acquired in a stationary system.

**Fig. 5.**
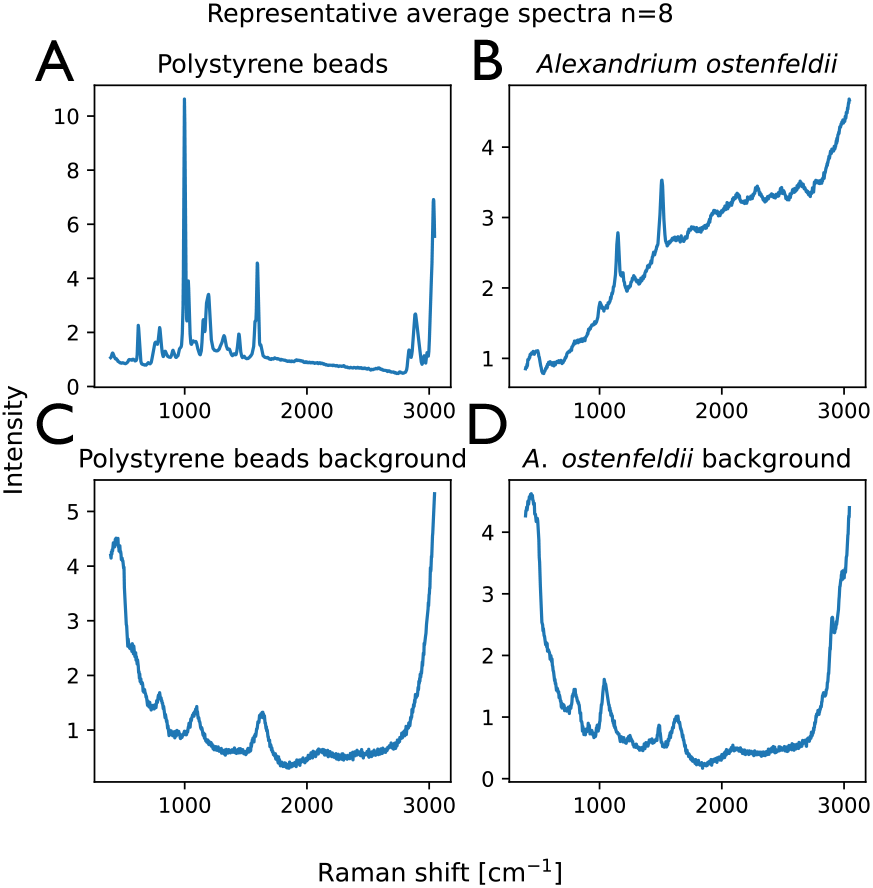
An average representative spectrum (n=8) for polystyrene beads and buffer background 3000 ppm HA suspended in MilliQ water (C) and *Alexandrium ostenfeldii*, a dinoflagellate algae (B) and 3000 ppm HA in 3 % instant ocean (D).

Here, we considered a positive detection as one that resulted in acquisition of a PS bead spectrum. On the other hand, a triggered detection that did not acquire a PS spectrum, but only showed background was categorized as false positive. An incoming particle that was detected by the speed measuring lasers, but did not yield a trigger event was a false negative. This could occur if it was too close to a preceding particle (see Supplementary Fig. S9) or if it was impossible to estimate its speed, for example due to a large deviation from the central alignment and thus a low speed. The proportion of false negatives increased linearly with the frequency of incoming particles (Fig. S10).

The system performance was tested using polystyrene beads comparing the number of issued trigger signals with the number of captured spectra (Fig. 6). The exposure time for the Raman measurement was set to 50 ms and the whole acquisition cycle was 200 ms. With these parameters, the system can collect a maximum of 80 spectra per minute (see blue line in Fig. 6). Previously reported similar studies achieved 4-8 spectra/min ^12^, 2 spectra/min ^45^ or 30 spectra/min ^20^, importantly these studies also implemented cell sorting, which might have influenced their throughput. Lyu *et al*. ^16^ show throughput of 300 spectra/min. However, their setup only focuses on single band at 1521 cm^−1^ of total width of 90 cm^−1^. Wang *et al*. ^15^ have shown superior throughput of 260 spectra/min, but their system does not contain any trigger mechanism and therefore is incompatible with many Raman detectors due to data shifting downtime. The highest reported throughput has been 3000 spectra/min combining spontaneous Raman with dielectrophoresis ^13^. Such system requires an additional dielectrophoresis trapping setup to achieve high performance, placing constraints on buffer conductivity and thus limiting its application.

**Fig. 6.**
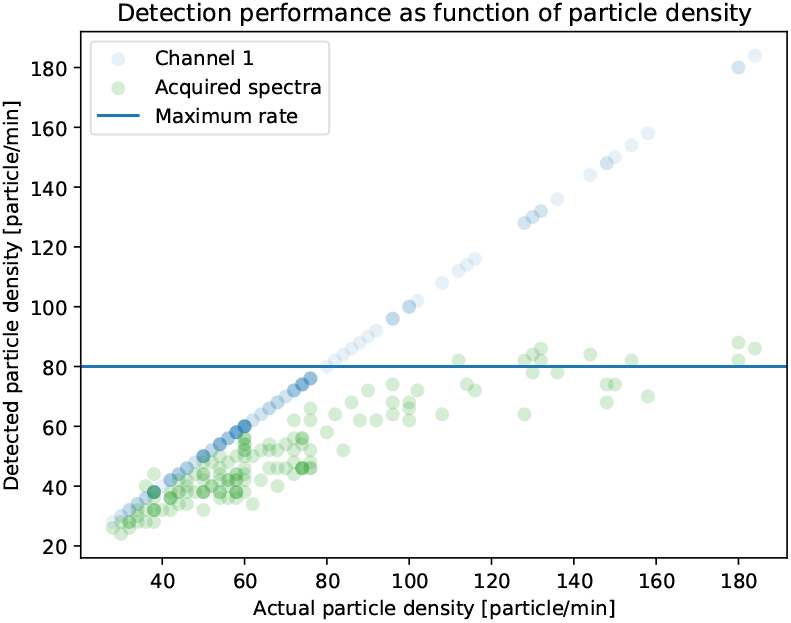
The rate of incoming PS particles per minute as detected by the ch1 laser is shown on the x axis. The data from the ch1 is also presented for clarity with blue circles. The y axis displays the number of Raman measurements acquired in the same window of time. The number of acquired Raman spectra is displayed with green circles. Maximum rate of detection for the tested camera is shown with a horizontal blue line.

Data was collected from four separate experiments with an average rate of false positive measurement of 1.91 % (SD = 0.35 %) (see Fig. S11). For individual false positive rates and number of particles in each experiment, see Supplementary table S1. This is a performance comparable to that of other systems ^13,15,20^

Due to the random distribution of the particles, some arrive at the detector within less than 200 ms from each other and thus are undetectable in a system without a particle capture system. The proportion of particles that were too close to be detected was generally between 10 and 35 % “for reasonable” particle density of 60 to 140 particle/min (see Fig S4, S5). The throughput could be further improved on acquisition devices with a faster acquisition loop, such as a detector with faster acquisition or lower exposure time. An additional challenge is that the number of slots reserved for driving GPIO signal in RPi5 is only 4. This means that the trigger scheduling system can get overwhelmed at high particle densities, which indeed was observed in our experiments. This can be overcome by using multiple trigger devices, which is easy to implement within our modular architecture and shared centralized timing.

### 3.6 Experiment with biological sample

We demonstrated the capacity of the setup to detect distinct biological spectra using the dinoflagellate alga *Alexandrium ostenfeldii*. In this experiment, we collected a total of 667 spectra, which were plotted using PCA to show distinct groupings of algal spectra, false negative detections and an unknown group of signals, possibly originating from empty dinoflagellate shells (Fig. 7). The alga spectrum is dominated by autofluorescence with resonant peaks at 1525 cm^−1^ and 1160 cm^−1^ see Fig. 5C and Fig 7D. Non-resonant Raman features are not resolved due to the overwhelming fluorescence. This shows the system capacity to acquire fully resolved spontaneous Raman spectra of flowing biological samples.

**Fig. 7.**
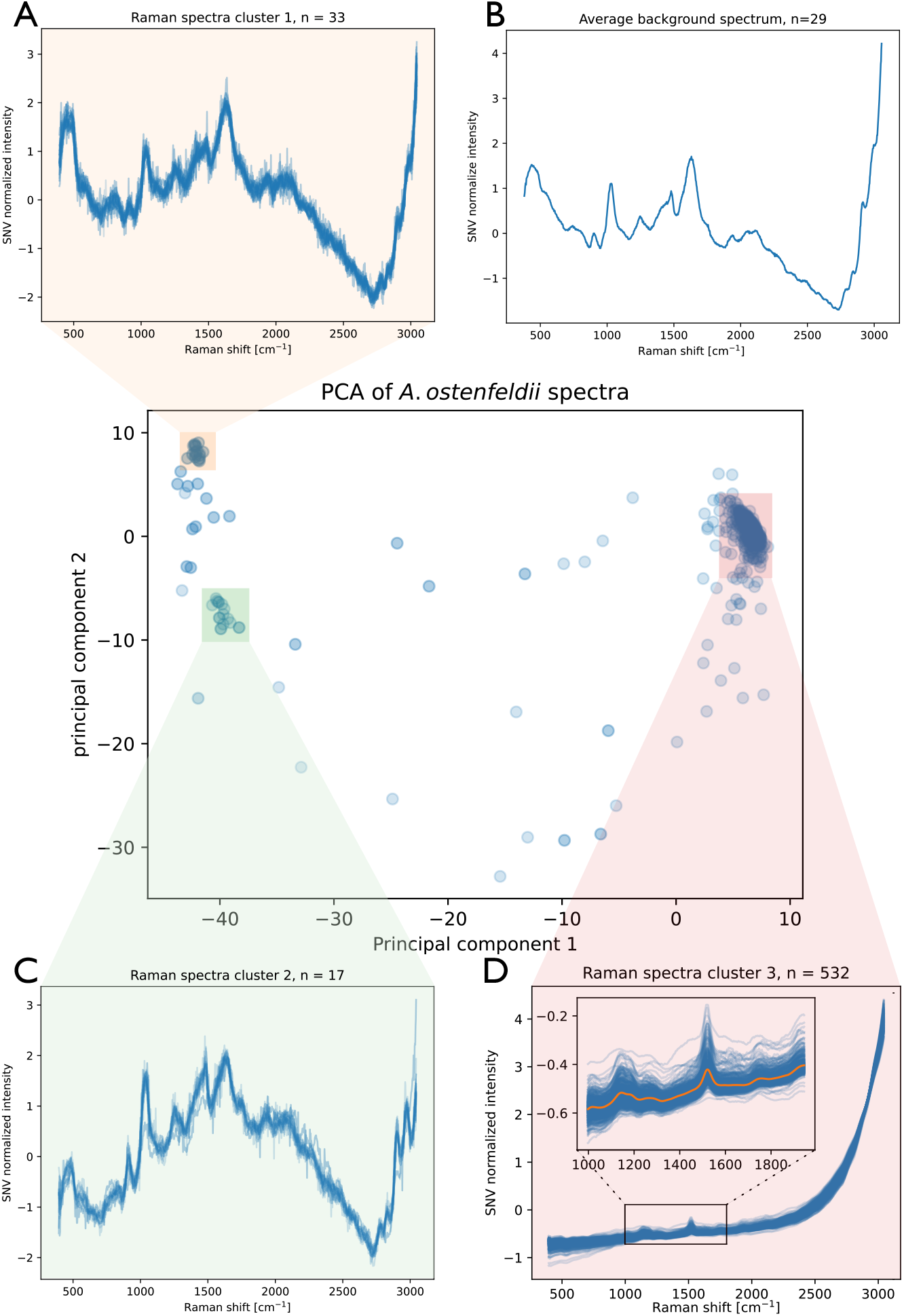
A principal component analysis of the *A. ostenfeldii* flow detection experiment. A total of 667 observations are presented and arbitrary areas are highlighted around the main clusters of data points. (A) Corresponds to false positives (empty background) as verified by (B) background controls acquired in the same setup. (C) A cluster of signals of unknown origin, possibly from empty dead dinoflagellate husks and (D) the surveyed *A. ostenfeldii* signal.

and We present an automated microfluidic platform capable of detecting incoming particles and precise triggering of a Raman detector (Fig 2). This platform directs biological sample through a fused silica capillary using a single pump, with the help of viscoelastic flow to passively focus particles in the center of the capillary. The particles are first detected by a pair of forward scatter detectors (channel 1 and 2; Fig. 2A) upon which the particle speed and time of arrival to the detector are calculated and a trigger signal is scheduled and actualized (Fig. 2C). The system employs a bespoke algorithm to predict the future position of incoming particles based on their velocity, enabling temporal coordination for downstream manipulations. A graphical user interface was implemented to facilitate control over flow operations and spectral data acquisition. The modular design of both hardware and software components allows for straightforward integration of additional functionalities. System performance has been validated using polystyrene beads and algal cells, demonstrating reliable detection and operational robustness.

Specific system parameters such as detector speed strongly influence the device throughput. Commercially available faster detectors would allow both for higher throughput and a reduced number of false negatives. However, the application relies on relatively high contrast of particles interacting with the speed detection lasers, which might make some samples difficult to study. Furthermore, this method relies on flow focusing, which might be challenging for highly heterogeneous samples, where diverse particulate shapes may affect flow speeds throughout the channel.

In contrast to previously published systems using polymeric microfluidic designs (e.g. PDMS ^15,16,20,21^ or PMMA ^46^), we present a system using only a fused silica capillary allowing for lower Raman background. Furthermore our setup uses easily replaceable off-the-shelf components using open source and a license free software system. Together with a plug-in based development approach, this makes the setup easily extendable and more widely accessible.

## Author contributions

MS, BFS, AK and AD contributed conceptualization and funding acquisition as well as project administration. AD, MBT and BFS provided microbiology related resources and supervision. AK provided optofluidic resources and supervision and SSK provided 3D printing resources and supervision. AS and MS wrote the original draft of the manuscript and provided data visualization. MBT, AK, AD, SSK and BFS provided manuscript review and editing. MS and BR provided methodology and investigation related to 3D printing. MS contributed to fluidics and optical methodology and investigation and AS contributed microbiological investigation and methodology. AS designed and implemented the software and hardware solution. AS and MS validated the setup.

## Data availability

Data is available at the https://doi.org/10.11583/DTU.31049842.

## Acknowledgements

This work was supported by a Novo Nordisk Foundation Exploratory Interdisciplinary Synergy Programme Grant, Combining Optics and Microfluidics to Identify, Isolate and Cultivate High-value Microbes (COMI-CULT), Grant No: NNF21OC0070888 awarded to BFS and AK. B Rezaei and S. S. Keller acknowledge financial support by the European Research Council under the Horizon 2020 Framework Programme Grant No: 772370PHOENEEX. For device fabrication, the authors acknowledge PolyFabLab, a research infrastructure financially supported by the Novo Nordisk Foundation, grant number NNF21OC0068814. We would like to acknowledge the generous gift of alga *A. ostenfeldii* from Prof. Per Juel Hansen, University of Copenhagen. We would also like to acknowledge Doron Rosenthal’s help with figure graphical design.

