## Supplementary File for "Viscoelastic flow-integrated Raman spectroscopy platform^†^"

### 1 Figure S1

To study the height-based variations we used varying aspect ratio (AR) in designs for the alignment guides (AR: height/width). Initial calibration was performed for the capillary by varying AR from 1 to 2. It was observed that AR in the range of 1.35 to 1.45 resulted in nearly square cross-sections after the fabrication, as shown in Figure S1.

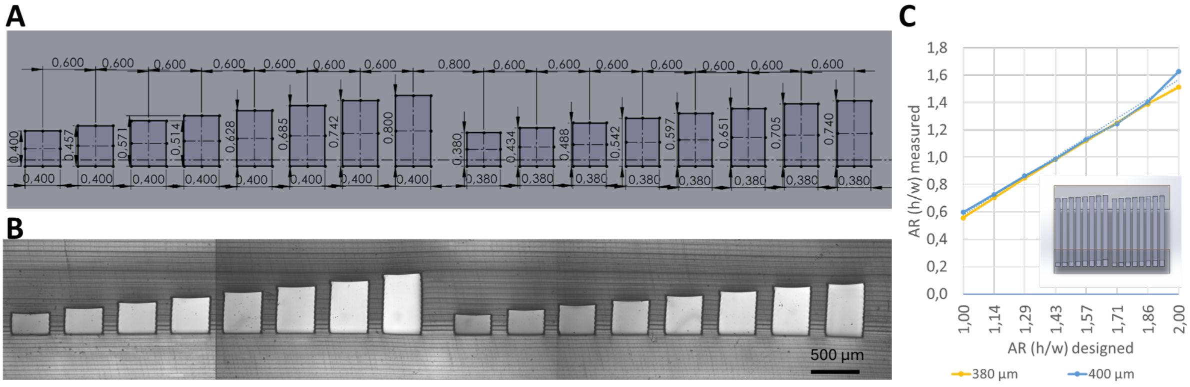

Figure 1: Square cross section optimization steps for fused silica capillary guides with widths of 400 and 380 µm. Top left is the CAD drawing (A), top right is the optical microscope images on the side wall of the printed part (C), and the right image is the comparison of AR for designed vs measured values after the printing and fabrication (B).

### 2 Figure S2

Figure S2 shows the calibration steps for guides of optical fibers with high AR. Width selected as 130 and 135 µm. Height was changed to get AR of 1.8 to 2.1. We observed approx. 25% downsizing at the height of the guides and prepared new set of experiments in Figure S3 to test AR of 1.4-1.8 for optical fiber and 1.25-1.35 for capillary alignment guides. Capillary guides tested for widths of 360, 365, 370, and 375 µm.

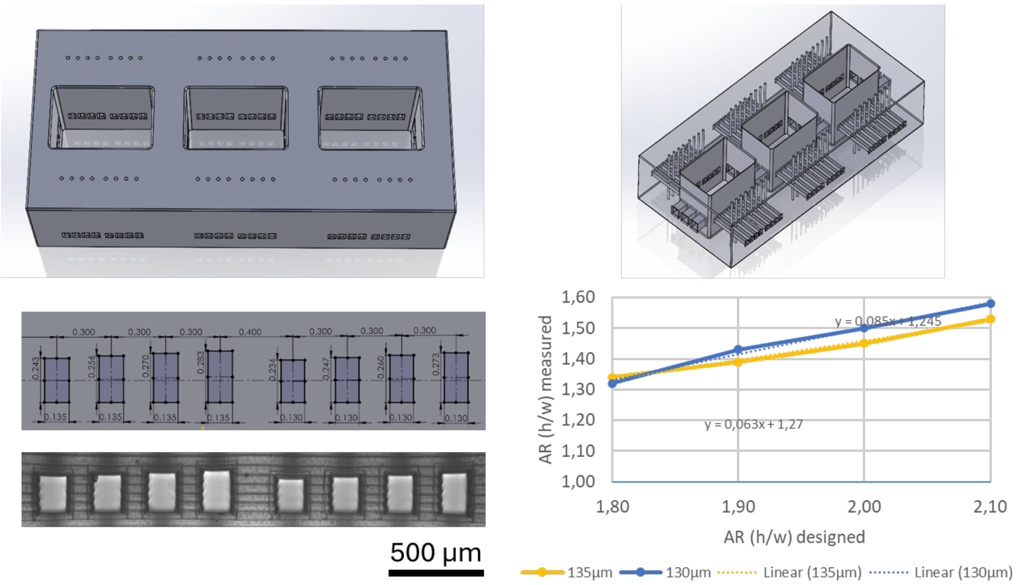

Figure 2: Square cross section optimization for fused optical fiber guides for large AR.

#### 3 Figure S3

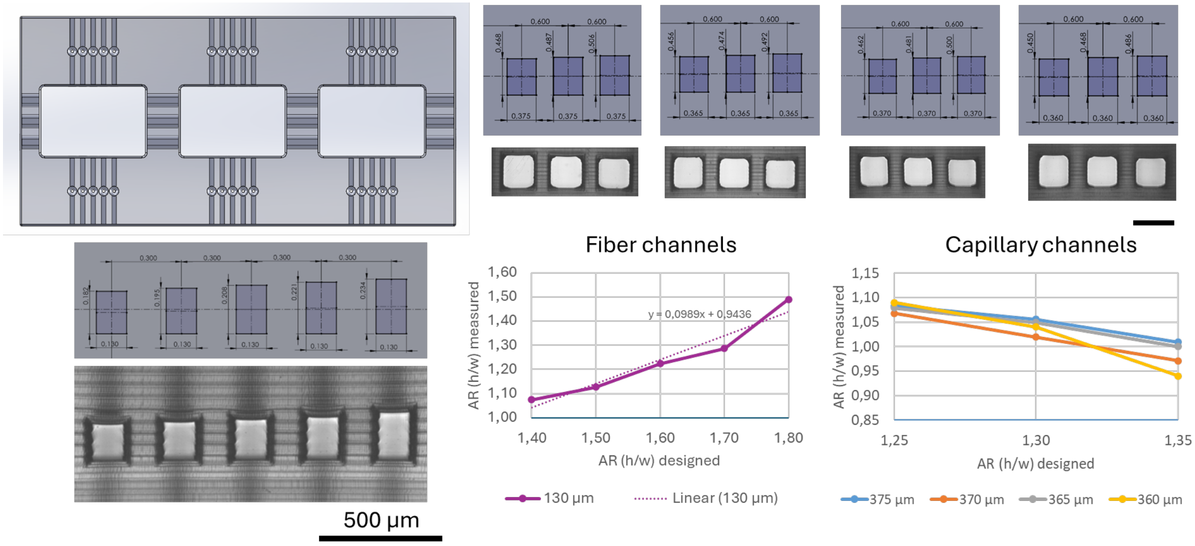

Figure 3: Square cross section optimization for optical fiber guides of AR: 1.4-1.8 and for silica capillary of AR: 1.25-1.35

### 4 Figure S4

Finally, in Figure S4 we decided to fabricate 5 guides for optical fibers with AR 1.3-1.5 and width of 127 and 130 µm and 3 guides for the capillary with AR 1.25-1.35 and width of 400 µm. Prior to finalizing the setup, the best fits were tested for each guide for optical fibers and capillary since their sizes from the shelf have variations.

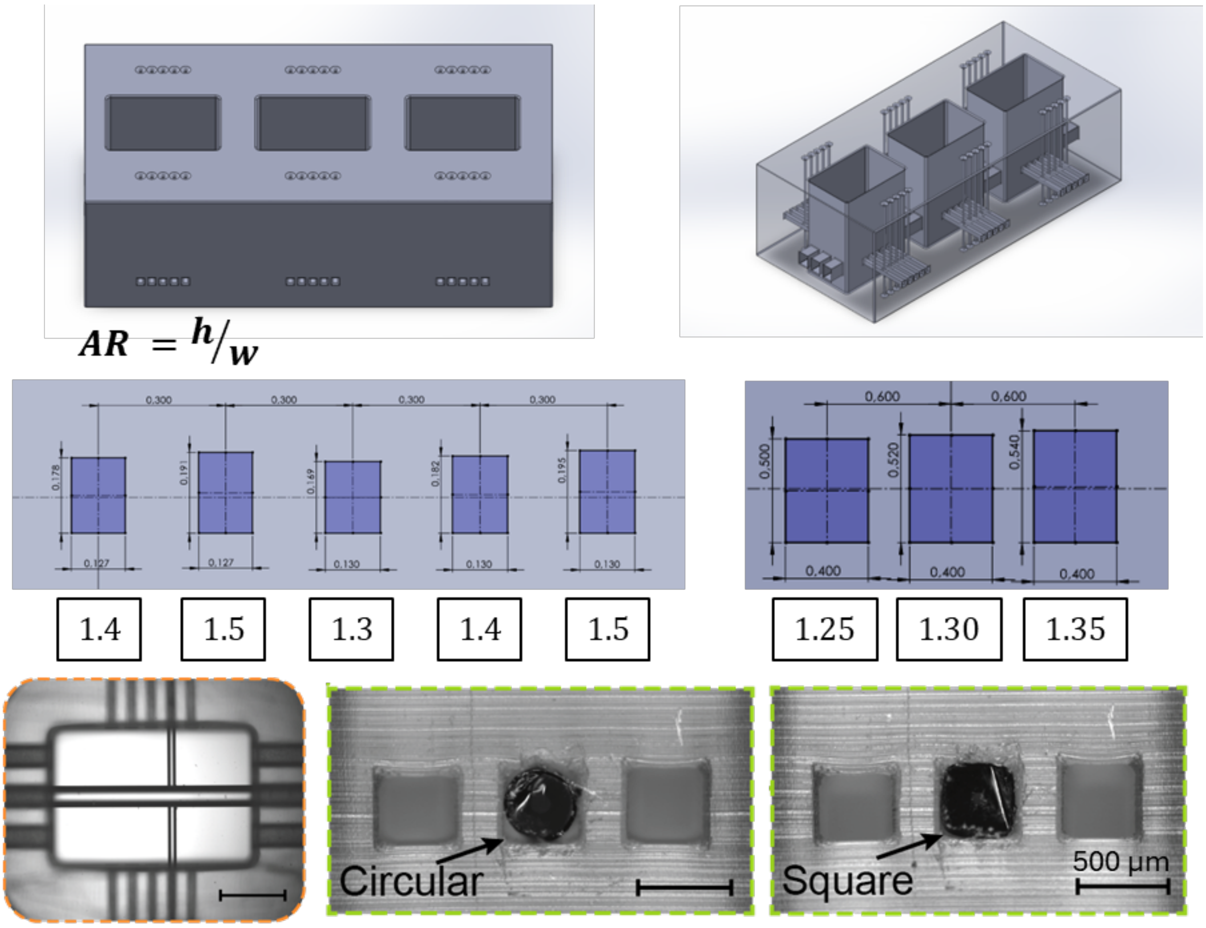

Figure 4: Final design demonstrates tight fit with optical fibers and fused silica capillaries at AR= 1.4 and 1.3, respectively.

### 5 Figure S5

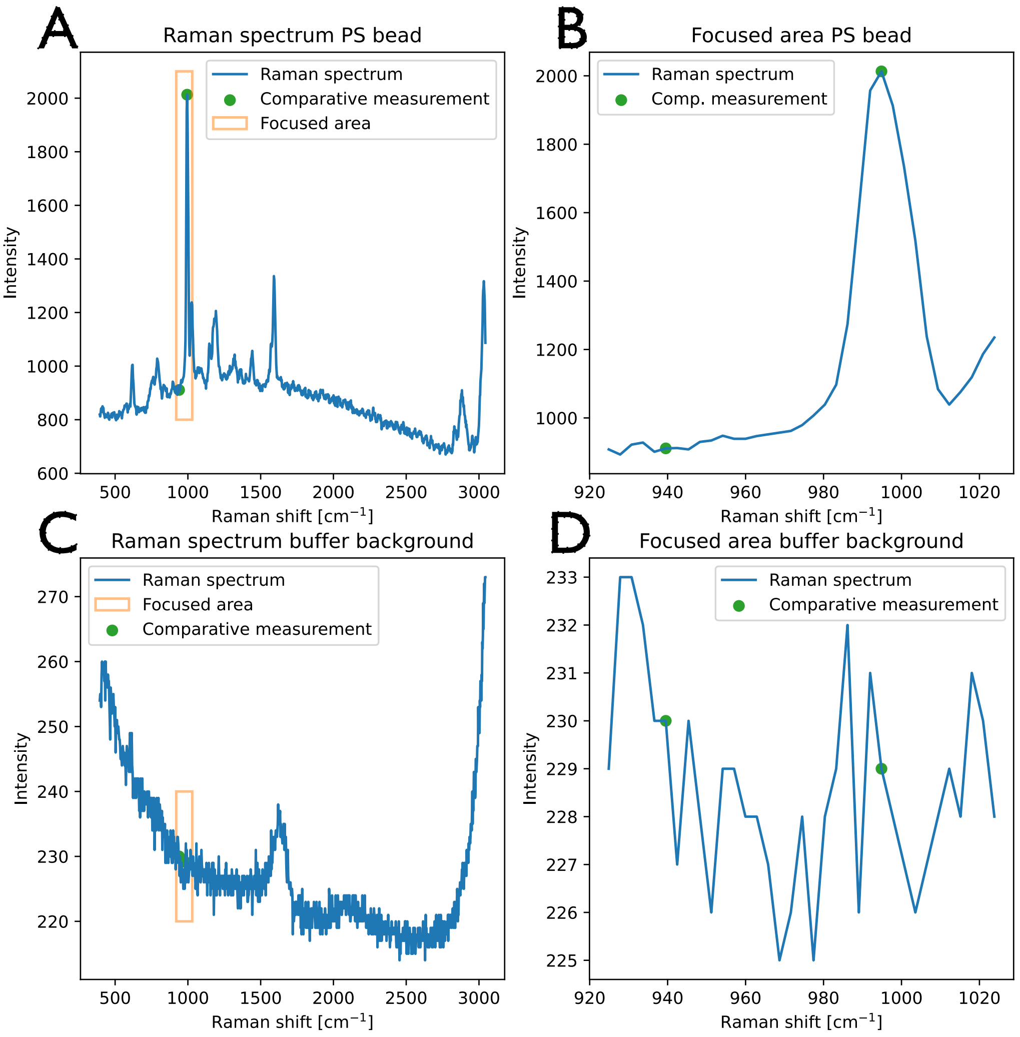

Figure 5: Detailed description of the process of selection of spectral features to identify false positive acquisitions. The highest peak wavelength (995.48 cm$^{-1}$) was compared to the low intensity region right next to this peak (A and B). This was contrasted with the background spectrum (C) where the same two spectral positions show similar intensity (D). A difference between those two spectral positions positions was calculated and used as a threshold value for false positive detection quantification.

#### 6 Figure S6

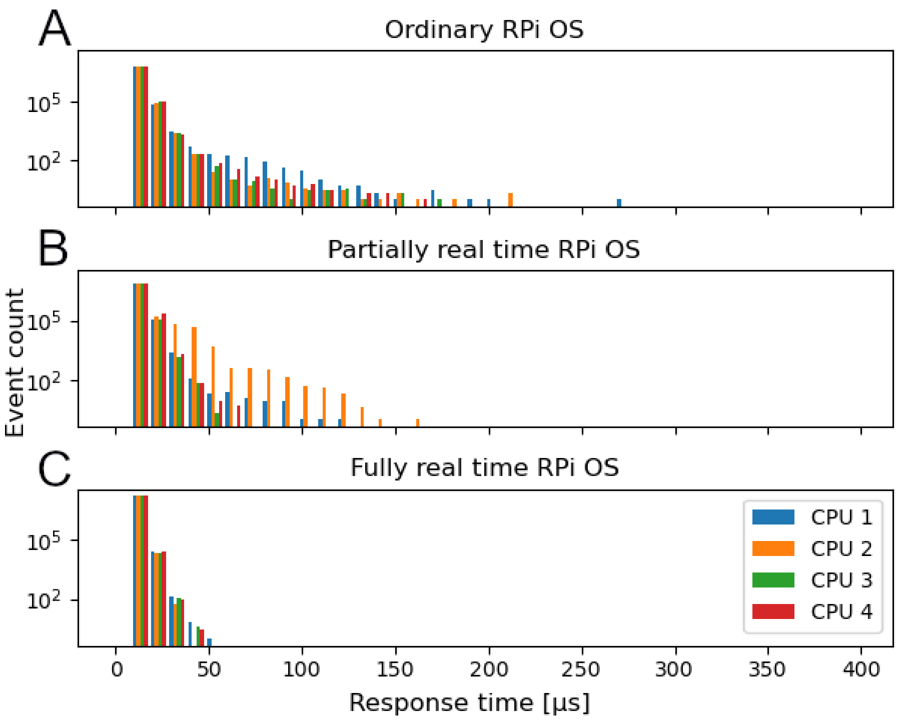

Figure 6: System latency under load comparison of an ordinary Raspberry Pi kernel (A), a partially RT-PREEMPT kernel (B) and fully RT-PREEMPT kernel. The x axis shows a task latency while the the y axis displays a logarithmic scale of the number of events recorded with a given latency.

#### 7 Figure S7

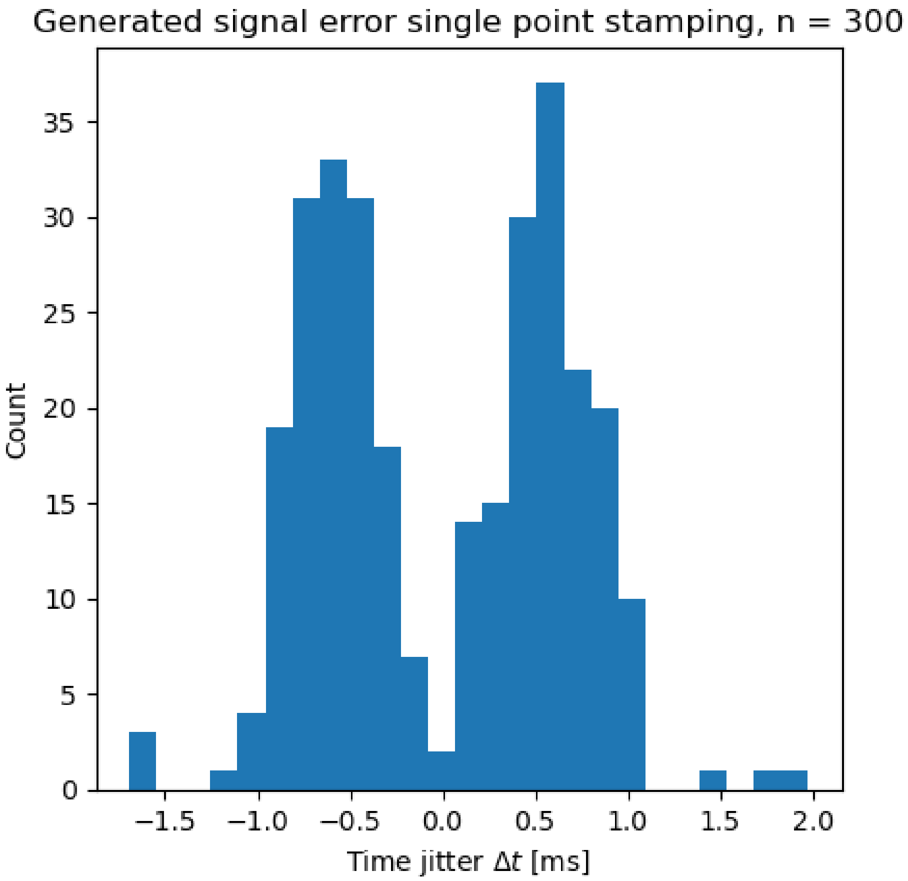

Figure 7: Error in the system time stamping of a generated square wave signal. A time difference from the real signal is shown in the x axis, while the y axis shows histogram counts for n = 300.

#### 8 Figure S8

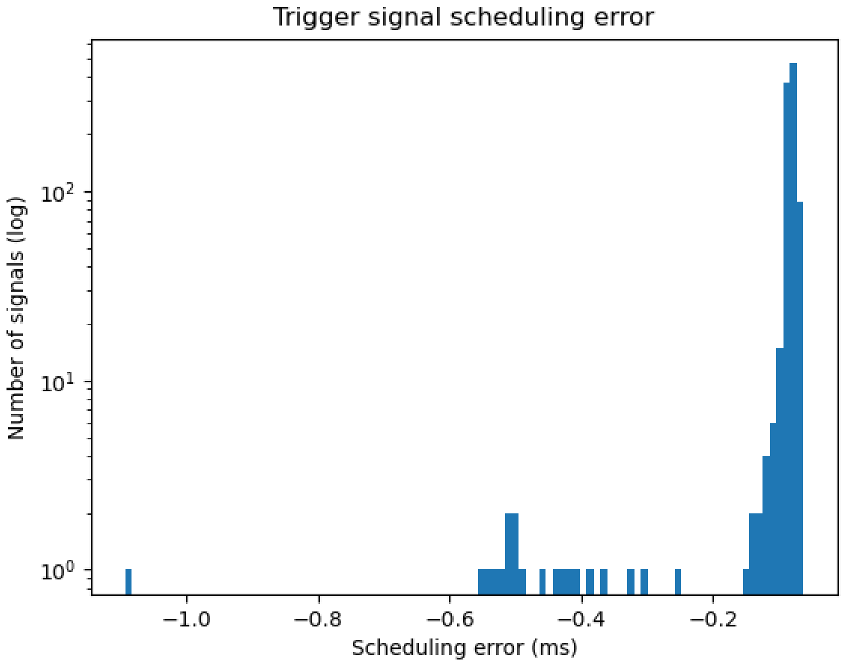

Figure 8: Histogram of the measured trigger signal scheduling error.

### 9 Figure S9

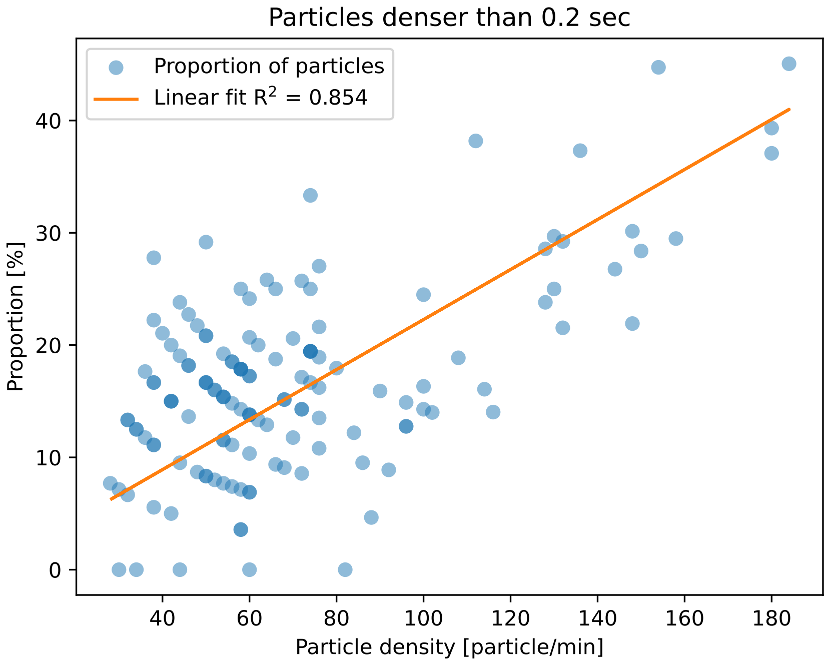

Figure 9: The y axis shows a proportion of particles that followed a previous particle in less than 0.2 sec, which is a timing limit of detection for our system. The x axis shows a total particle density in a given window of measurement. Total number of measurements is n = 141. An ordinary least squares regression was applied with p value $< 0.001$.

#### 10 Figure S10

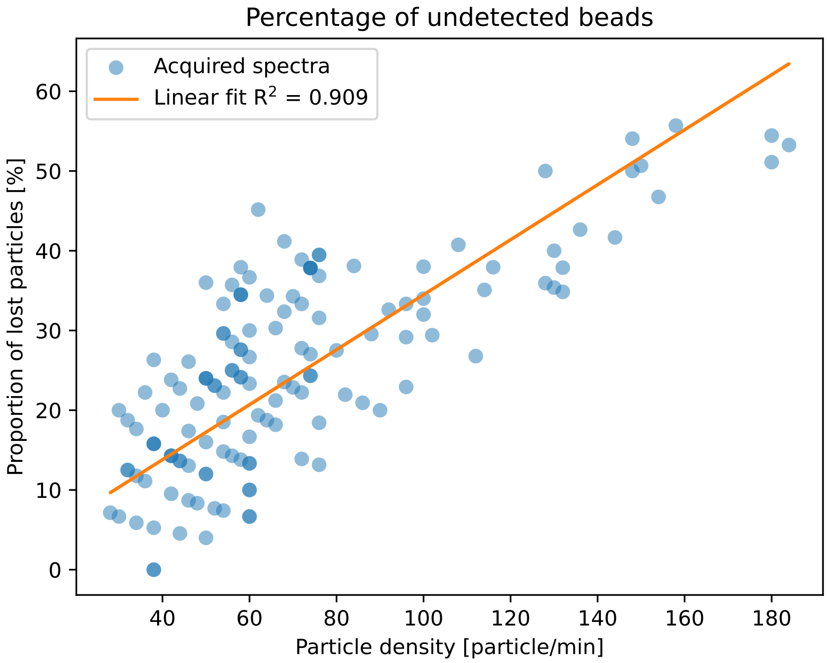

Figure 10: Proportion of particles that appeared in the channel 1, but did not yield a detection event (y axis) compared to a number of particles in a given measurement interval (y axis) n = 141. An ordinary least squares regression was applied with p value $< 0.001$.

#### 11 Figure S11

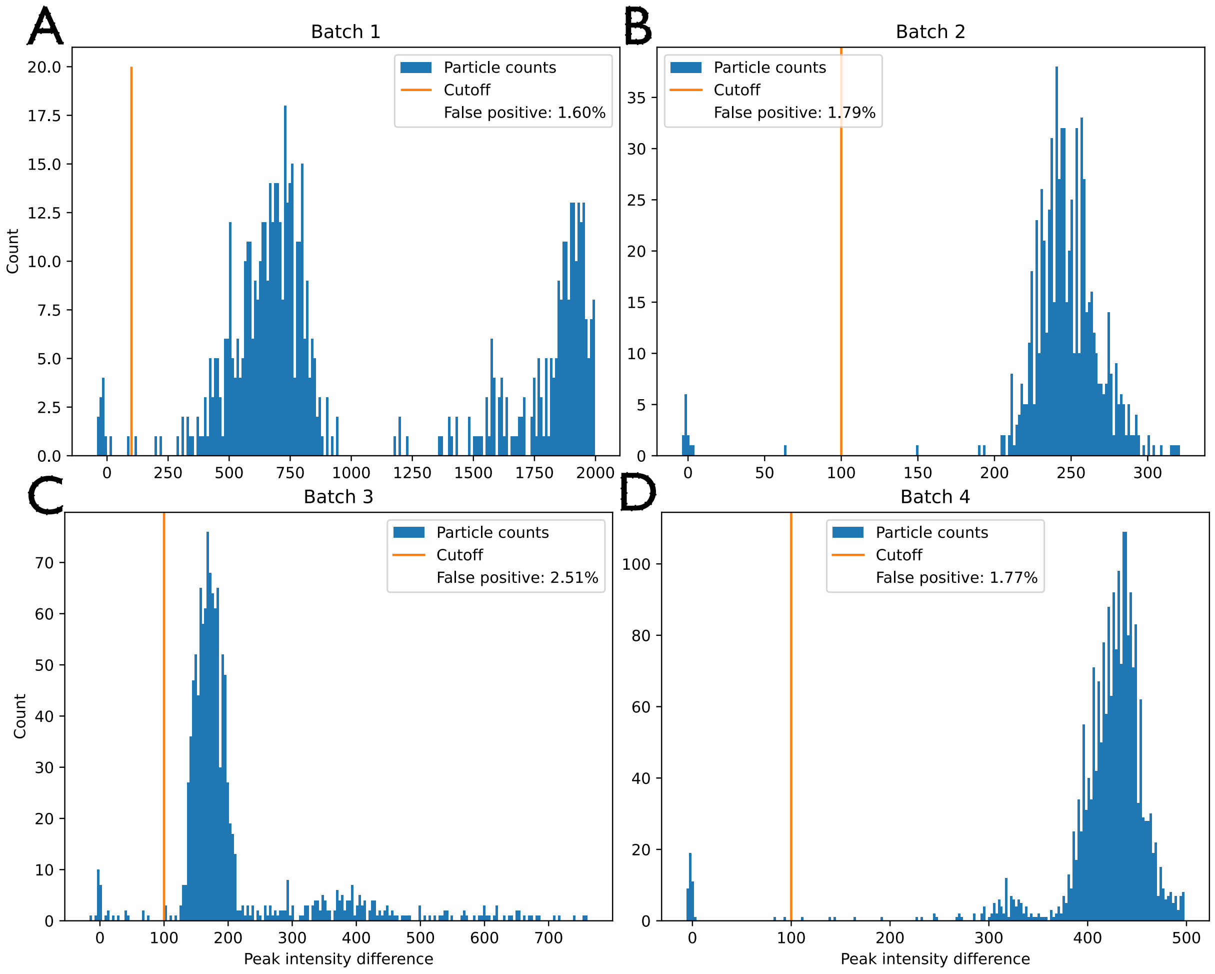

Figure 11: Comparison of the intensity of the differnce between the largest peak in the PS spectrum and a flat region next to it. Data was acquired over 4 runs over separate days to ensure stability and it is plotted as a histogram of the diffence between the highest peak and the valley. The selected cutoff was 100 for each measurement batch, shown with an orange line.

#### 12 Table S1

| Pressure [mBar] | Channel position | Average position [µm] |
| --- | --- | --- |
| 100 | Beginning | $144.95 \pm 7.92$ |
| | Raman detection area | $151.38 \pm 0.29$ |
| 500 | Beginning | $146.35 \pm 1.22$ |
| | Raman detection area | $143.00 \pm 9.09$ |
| 900 | Beginning | $150.39 \pm 7.63$ |
| | Raman detection area | $150.76 \pm 0.42$ |

Table 1: Average sppeds of particles measured at the beginning of the capillary (2 cm from the entrance) and within the Raman detection zone (20 cm from the entrance). Values are presented with a standard deviation.

### 13 Setting up RT-PEEMPT on Raspberry Pi

#### 13.1 Check or modify the configuration

Following a previously published manual the following steps should be performed to verify the kernel compilation.

1. Installing dependencies for building.

```
sudo apt install git bc bison flex libssl-dev make
```

2. Downloading linux kernel.

```
git clone --depth=1 https://github.com/raspberrypi/linux
```

3. Run the following command in the folder with the downloaded linux environment:

```
~/rpi-kernel/linux/$ make menuconfig
```

   - If you get the error make menuconfig' requires the `ncurses` libraries, install the `ncurses` library using:

   ```
   ~/rpi-kernel/linux/$ sudo apt-get install libncurses-dev
   ```

4. The most important configurations are:

   - Enable `CONFIG\_PREEMPT\_RT\_FULL`: General setup → Preemption Model (Fully Preemptible Kernel (RT)) → Fully Preemptible Kernel (RT)
   - Enable `HIGH\_RES\_TIMERS`: General setup → Timers subsystem → High Resolution Timer Support
   - Set `CONFIG\_HZ` to 1000Hz (read the notes!): Kernel Features → Timer frequency = 1000 Hz

For details refer to the linked tutorial above.

#### 13.2 Kernel compilation

This procedure generally follows the official Raspberry Pi manual.

1. Downloading the RT_PREEMPT patch. `https://mirrors.edge.kernel.org/pub/linux/kernel/projects/rt/6.6/` NOTE: Make sure the kernel version corresponds to the RPi kernel.
2. Unzip patch and copy it to the linux kernel repository folder.
3. Run the patch

```
cat patch-6.6.25-rt29.patch | patch -p1
```

4. Configure the kernel compilation

```
KERNEL=kernel_2712
make bcm2712_defconfig
```

5. Before the next step, one might want to modify the configuration file (`/boot/firmware/conf.txt`).
6. Edit the line to give the kernel a specific name.

```
CONFIG_LOCALVERSION="-v7l-MY_CUSTOM_KERNEL"
```

7. Build kernel by running:

```
make -j4 Image.gz modules dtbs
sudo make modules_install
sudo cp arch/arm64/boot/dts/broadcom/*.dtb /boot/firmware/
sudo cp arch/arm64/boot/dts/overlays/*.dtb* /boot/firmware/overlays/
sudo cp arch/arm64/boot/dts/overlays/README /boot/firmware/overlays/
sudo cp arch/arm64/boot/Image.gz /boot/firmware/$KERNEL.img
```

8. Edit the `config.txt` file to select the kernel that the Raspberry Pi will boot.

```
kernel=kernel-myconfig.img
```

#### 13.3 Checking the performance

This broadly follows previously mentioned LeMaRiva method.

The suite needs the `build-essential` package in order to be compiled in Raspbian/Debian. To install the RT-Test suite follow these steps:

```
sudo apt-get install -y build-essential
# if you are using the Raspbian lite,
# you need to install git: sudo apt-get install -y git
git clone git://git.kernel.org/pub/scm/utils/rt-tests/rt-tests.git
cd rt-tests
git checkout stable/v1.0
make all -j4
sudo make install
```

You need to check out the `stable/v1.0` branch to build the project. If you want to only compile the `cyclictest`, just replace `make all -j4` with make `cyclictest`.

##### 13.3.1 Running RT-Test

This test only generates data that are subsequently plotted in python. The `cyclictest` was run with the following parameters:

```
sudo cyclictest -l50000000 -m -S -p90 -i200 -h400 -q > output.txt
```

Where individual settings are as follows:

`-l50000000`: 50M iterations (about 2.5 hours!);
`-m`: lock current and future memory allocations (prevent being paged out?)
`-S`: Standard SMP testing: options -a -t -n and same priority of all threads (Raspberry Pi has 4 Cores then 4 Threads)
`-p90`: priority of highest prio thread set to 90 (for the 4 threads, then: 90 89 88 87)
`-i200`: interval for the first thread (in us).
`-h1000`: dump histogram for max latency (up to 1000us).
`-q`: print a summary only on exit.

Test will run indefinitely until it is interrupted by the user.

A load is simulated using `hackbench` which is a part of the RT-Test. This needs to be run as `sudo` (super user) otherwise it will result in an error: `unable to change scheduling policy`

After the test concludes, you get the file `output.txt` with the results. Typing the following:

```
grep -v -e "^#" -e "^$" output.txt | tr " " "," | tr "\t" "," > histogram.csv
sed -i '1s/^/time,core1,core2,core3,core4\n /' histogram.csv
```

This retrieves the data lines, removes empty lines and creates a comma-separated values file (.csv).

This data can be used with Python to make some analysis and plots.

#### 13.4 Test results

##### 13.4.1 Regular kernel

```
uname -a
Linux test 6.6.20+rpt-rpi-2712 #1 SMP PREEMPT Debian 1:6.6.20-1+rpt1
    (2024-03-07) aarch64 GNU/Linux

hackbench -l 500000
Running in process mode with 10 groups using 40 file descriptors each
    (== 400 tasks)
Each sender will pass 500000 messages of 100 bytes
Time: 1046.860

sudo cyclictest -l50000000 -m -S -p90 -i200 -h400 -q >
    ordinaryKernel_cyclictest3withload.txt
```

Notes: Temperature of the CPU went up to 60°C.

##### 13.4.2 RT kernel with intermediate preemption

This was an out of the box patch with no settings in the `config.txt` changed.

```
uname -a
Linux test 6.6.26-rt29-v8-16k-MY_RT+ #1 SMP PREEMPT Mon Apr 15 14:55:43
     CEST 2024 aarch64 GNU/Linux

sudo cyclictest -l50000000 -m -S -p90 -i200 -h400 -q >
    RT1partpreempt_cyclictest.txt
```

##### 13.4.3 Fully preemptible RT kernel

```
uname -a
Linux test 6.6.26-rt29-v8-16k-MY_FULLY_RT+ #1 SMP PREEMPT_RT Mon Apr 15
     18:35:03 CEST 2024 aarch64 GNU/Linux

sudo cyclictest -l50000000 -m -S -p90 -i200 -h400 -q >
    RT2fullyPreemt_cyclictest1.txt

hackbench -l 500000
Running in process mode with 10 groups using 40 file descriptors each
    (== 400 tasks)
Each sender will pass 500000 messages of 100 bytes
Time: 2794.754
```

##### 13.4.4 Grandmaster fully preemptible RT kernel

```
uname -a
Linux grandmaster 6.6.32-rt32-v8-16k_fullRT_grandmaster-g5882bc5db17d-
    dirty #1 MP PREEMPT_RT Fri Jun  7 14:53:05 CEST 2024 aarch64 GNU/
    Linux

hackbench -l 500000
Running in process mode with 10 groups using 40 file descriptors each
    (== 400 tasks)
Each sender will pass 500000 messages of 100 bytes
```
